# Enzyme promiscuity shapes evolutionary innovation and optimization

**DOI:** 10.1101/310946

**Authors:** Gabriela I. Guzmán, Troy E. Sandberg, Ryan A. LaCroix, Akos Nyerges, Henrietta Papp, Markus de Raad, Zachary A. King, Trent R. Northen, Richard A. Notebaart, Csaba Pál, Bernhard O. Palsson, Balázs Papp, Adam M. Feist

**Affiliations:** Department of Bioengineering, University of California, San Diego, La Jolla, CA, 92093, United States.; Synthetic and Systems Biology Unit, Institute of Biochemistry, Biological Research Centre of the Hungarian Academy of Sciences, Szeged, Hungary H-6726.; Environmental Genomics and Systems Biology Division, Lawrence Berkeley National Laboratory, Berkeley, CA 94720, United States.; Laboratory of Food Microbiology, Wageningen University and Research, Wageningen, The Netherlands.; Novo Nordisk Foundation Center for Biosustainability, Technical University of Denmark, Lyngby, Denmark.; Department of Pediatrics, University of California, San Diego, La Jolla, CA, 92093, United States.

**Keywords:** adaptive evolution, enzyme promiscuity, systems biology, genome-scale modeling

## Abstract

Evidence suggests that novel enzyme functions evolved from low-level promiscuous activities in ancestral enzymes. Yet, the evolutionary dynamics and physiological mechanisms of how such side activities contribute to systems-level adaptations are poorly understood. Furthermore, it remains untested whether knowledge of an organism’s promiscuous reaction set (‘underground metabolism’) can aid in forecasting the genetic basis of metabolic adaptations. Here, we employ a computational model of underground metabolism and laboratory evolution experiments to examine the role of enzyme promiscuity in the acquisition and optimization of growth on predicted non-native substrates in *E. coli* K-12 MG1655. After as few as 20 generations, the evolving populations repeatedly acquired the capacity to grow on five predicted novel substrates–D-lyxose, D-2-deoxyribose, D-arabinose, m-tartrate, and monomethyl succinate–none of which could support growth in wild-type cells. Promiscuous enzyme activities played key roles in multiple phases of adaptation. Altered promiscuous activities not only established novel high-efficiency pathways, but also suppressed undesirable metabolic routes. Further, structural mutations shifted enzyme substrate turnover rates towards the new substrate while retaining a preference for the primary substrate. Finally, genes underlying the phenotypic innovations were accurately predicted by genome-scale model simulations of metabolism with enzyme promiscuity. Computational approaches will be essential to synthesize the complex role of promiscuous activities in applied biotechnology and in models of evolutionary adaptation.

## 1. Introduction

Understanding how novel metabolic pathways arise during adaptation to environmental changes remains a central issue in evolutionary biology. The prevailing view is that enzymes often display promiscuous (i.e., side or secondary) activities and evolution takes advantage of such pre-existing weak activities to generate metabolic novelties[1, 2, 3, 4, 5, 6, 7, 8, 9]. However, it remains poorly explored how and at what evolutionary stages enzyme side activities contribute to environmental adaptations. Do genetic elements associated with promiscuous activities mutate mostly in the initial ‘innovation’ stage of adaptation when the population acquires the ability to grow on a new nutrient source[9, 10] (i.e., innovation) or do they also contribute to improving fitness in subsequent stages (i.e., optimization)[11]? Innovations have been linked to beneficial mutations that endow an organism with novel capabilities such as the ability to use a new carbon source and expand into a new ecological niche[11, 12]. This is distinct from optimizations associated with mutations that improve upon the initial innovation. It is often observed that the mutations accrued within this optimization phase produce gradual benefits in fitness[11]. Typically, enzyme promiscuity has been linked to the innovation phase, for which mutations enhancing secondary activities may result in dramatic phenotypic improvements[2, 11]. In this work, we demonstrate that enzyme promiscuity can be linked to fitness benefits in both the innovation and optimization stages of adaptive evolution.

A second open question in understanding the role of enzyme promiscuity in adaptation concerns our ability to predict the future evolution of broad genetic and phenotypic changes[13, 14]. There has been an increasing interest in studying empirical fitness landscapes to assess the predictability of evolutionary routes[15, 16]. However, these approaches assess predictability only in retrospect and there is a need for computational frameworks that forecast the specific genes that accumulate mutations based on mechanistic knowledge of the evolving trait. A recent study suggested that a detailed knowledge of an organisms promiscuous reaction set (the so-called ‘underground metabolism’[17]) enables the computational prediction of genes that confer new metabolic capabilities when artificially overexpressed[4]. However, it remains unclear whether this approach could predict evolution in a population of cells adapting to a new nutrient environment through spontaneous mutations. First, phenotypes conferred by artificial overexpression might not be accessible through single mutations arising spontaneously. Second, and more fundamentally, mutations in distinct genes may lead to the same phenotype.

Such alternative mutational trajectories may render genetic evolution largely unpredictable. Furthermore, computational approaches can aid in predicting and discovering overlapping physiological functions of enzymes [15, 18], but these have also yet to be explored in the context of adaptation. In this study, we address these issues by performing controlled laboratory evolution experiments to adapt *E. coli* to predicted novel carbon sources and by monitoring the temporal dynamics of adaptive mutations.

## 2. Results and Discussion

### 2.1. Computational prediction and experimental evolution of non-native carbon source utilizations

To test our ability to predict evolutionary adaptation to novel (non-native) carbon sources based on our knowledge of underground metabolism, we utilized a comprehensive network reconstruction of underground metabolism[4]. This network reconsruciton was previously shown to correctly predict growth on non-native carbon sources if a given enabling gene was artificially overexpressed in a growth screen[4]. By adding the set of underground reactions to the comprehensive metabolic reconstruction for *E. coli* K-12 MG1655, *i*JO1366[19], we employed the constraint-based modeling framework to identify carbon sources where the native *E. coli* metabolic network was unable to grow, but addition of a single underground reaction predicted growth. Based on this computational procedure, we selected eight carbon sources that cannot be utilized by wild-type *E. coli* MG1655 (Table S1).

Next, we initiated laboratory evolution experiments to adapt *E. coli* to these non-native carbon sources to examine the validity of the computational predictions. Adaptive laboratory evolution experiments were conducted in two distinct phases: first, an ‘innovation’[9, 10] stage during which cells acquired mutations to grow on the non-native carbon sources and, second, an ‘optimization’[11] stage during which a strong pressure was placed to select for the fastest growing cells on the novel carbon sources (Figure 1A).

**Figure 1:**
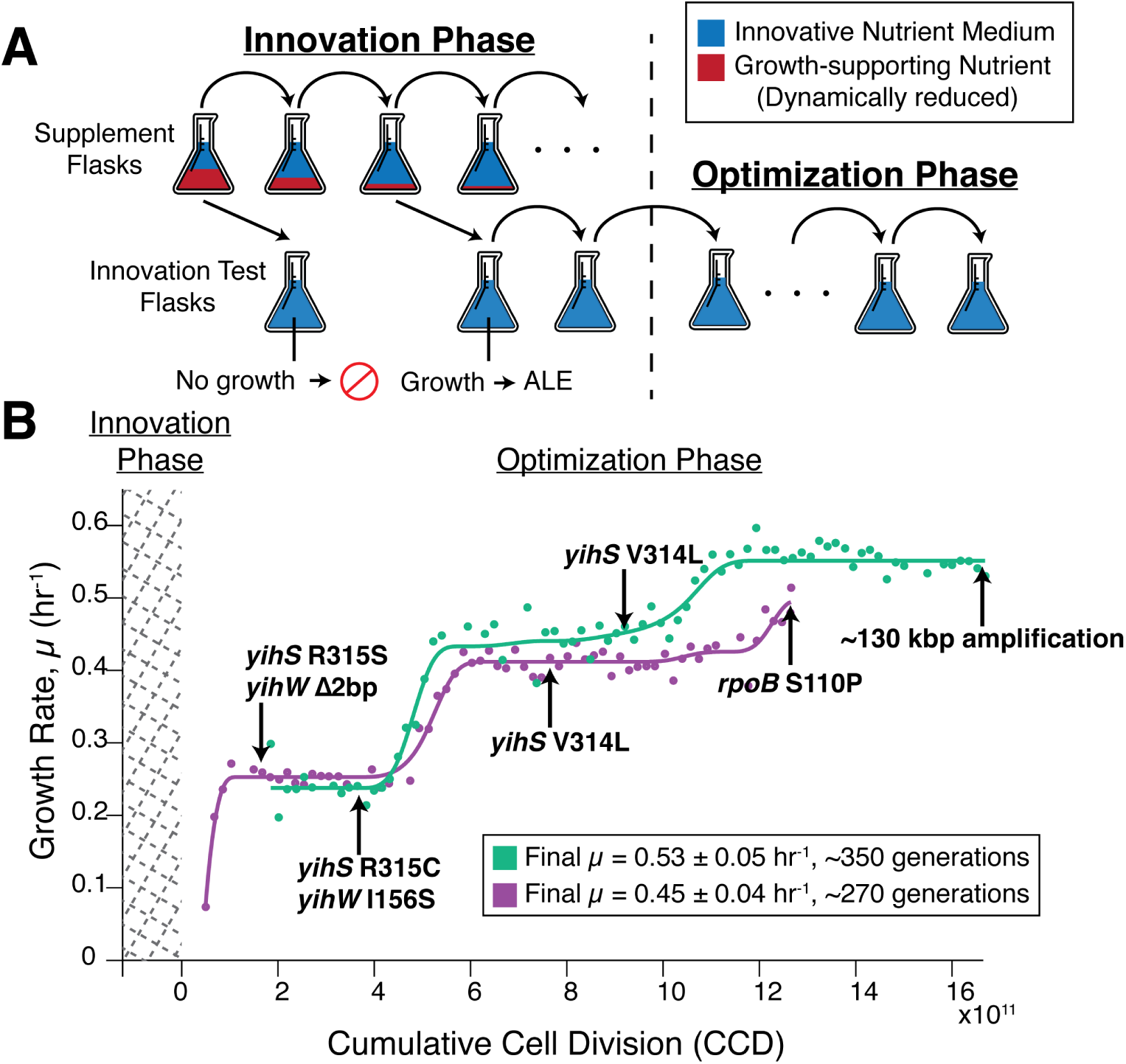
Laboratory evolution method schematic and the growth trajectory of D-lyxose experiments. A) A schematic of the two-part adaptive laboratory evolution (ALE) experiments. The innovation phase involved growing cells in supplemented flasks containing the innovative substrate (blue) and growth-promoting supplement (red). As cultures were serially passed, they were split into another supplemented flask as well as an ‘innovation test flask’ containing only the innovative nutrient (no supplement) to test for the desired evolved growth phenotype. The ‘optimization’ phase consisted of selecting for the fastest growing cells and passing in mid log phase. B) Growth rate trajectories for duplicate experiments (green and purple) for the example case of D-lyxose. Population growth rates are plotted against cumulative cell divisions. Clones were isolated for whole genome sequencing at notable growth-rate plateaus as indicated by the arrows. Mutations gained at each plateau are highlighted beside the arrows (mutations arising earlier along the trajectory persisted in later sequenced clones).

During the initial innovation stage of laboratory evolution experiments (Figure 1A, see *SI Materials and Methods*), *E. coli* was successfully adapted to grow on five non-native substrates. Duplicate laboratory evolution experiments were conducted in batch growth conditions and in parallel on an automated adaptive laboratory evolution (ALE) platform using a protocol that uniquely selected for adaptation to conditions where the ancestor (i.e., wild-type) was unable to grow (Fig. 1A)[20]. In the innovation phase, *E. coli* was weaned off of a growth-supporting nutrient (glycerol) onto the novel substrates (Fig. 1A, Table S2). A description of the complex passage protocol is given in the Figure 1 legend and expanded in the methods for both phases of the evolution. This procedure successfully adapted *E. coli* to grow on five out of eight non-native substrates, specifically, D-lyxose, D-2-deoxyribose, D-arabinose, m-tartrate, and monomethyl succinate. Unsuccessful cases could be attributed to various experimental and biological factors such as experimental duration limitations, the requirement of multiple mutation events, or stepwise adaptation events, as observed in an evolving *E. coli* to utilize ethylene glycol [21].

### 2.2. Underground metabolism accurately predicted the genes mutated during innovation

To analyze the mutations underlying the nutrient utilizations, clones were isolated and sequenced shortly after an innovative growth phenotype was achieved and mutations were identified (see Methods) and analyzed for their associated causality (Fig. 1B, Fig. S1, Dataset S1). Strong signs of parallel evolution were observed at the level of mutated genes in the replicate evolution experiments. Such parallelism provided evidence of the beneficial nature of the observed mutations and is a prerequisite for predicting the genetic basis of adaptation[22]. Mutations detected in the evolved isolated clones for each experiment demonstrated a striking agreement with such predicted ‘underground’ utilization pathways[4]. Specifically, for four out of the five different substrate conditions, key mutations were linked to the predicted enzyme with promiscuous activity, which would be highly unlikely by chance (P <10^-8^, Fishers exact test), (Table 1, Fig. S2). Not only were the specific genes (or their direct regulatory elements) mutated in four out of five cases, but few additional mutations (0-2 per strain, Dataset S1) were observed in the initial innovation phase, indicating that the innovations required a small number mutational steps to activate the predicted growth phenotype and the method utilized was highly selective. For the one case where the prediction and observed mutations did not align, D-arabinose, a detailed inspection of the literature revealed existing evidence that three *fuc* operon associated enzymes can metabolize D-arabinose–FucI, FucK, and FucA [23]. In this case, the modeling approach could not make the correct prediction because the promiscuous (underground) reaction database was incomplete.

**Table 1:** Key Innovative Mutations.

| Gene Mutated | Substrate | Gene Prediction | Protein Change(s) (Experiment #) | Perceived Impact (Structural (S) or Regulatory (R)) |
| --- | --- | --- | --- | --- |
| *yihS* | D-Lyxose | *yihS* | R315S(1) | Substrate binding<sup>1</sup> (S) |
|  |  |  | R315C(2) | Substrate binding<sup>1</sup> (S) |
| *yihW* | D-Lyxose | *yihS* | Frameshift(1) | Loss of function, large truncation (R) |
|  |  |  | I156S(2) | - (R) |
| *rbsK* | D-2-Deox. | *rbsK* | N20Y(1) | - (S) |
| *rbsR* | D-2-Deox. | *rbsK* | Insertion Sequence(1) | Loss of function, increased *rbsK* expression (R) |
| 181 kbp and 281 kbp Regions | D-2-Deox. | *rbsK* | Large Duplications(1) | Increased gene expression (R) |
|  |  |  | 165 and 262 genes |  |
| *fucR* | D-Arabinose | *rbsK* | D82Y(1) | Pfam: DeoRC C terminal substrate sensor domain<sup>2</sup> (R) |
|  |  |  | S75R(1 and 2) | Pfam: DeoRC C terminal substrate sensor domain<sup>2</sup> (R) |
|  |  |  | *244C(2) | - (R) |
| *dmlA* | m-Tartrate | *dmlA* | A242T(1) | - (S) |
| *dmlR/dmlA* | m-Tartrate | *dmlA* | intergenic -50/-53(2) | sigma 70 binding: -10 of dmlRp3 promoter<sup>3</sup> (R) |
|  |  |  | intergenic -35/-68(2) | dmlRp3 promoter region<sup>3</sup> (R) |
| *ybfF/seqA* | Mon. Succ. | *ybfF* | intergenic -73/-112(1) | sigma 24 binding: -35 of ybfFp1 promoter<sup>3</sup> (R) |
|  |  |  | intergenic -51/-123(2) | sigma 24 binding: -10 of ybfFp1 promoter<sup>3</sup> (R) |

Substrates D-2-deoxyribose and monomethyl succinate are abbreviated D-2-Deox. and Mon. Succ. respectively. The detailed locations of the mutations listed in this table are available in Datasets S1. <sup>1</sup>Substrate binding information about YihS previously published[34]. <sup>2</sup>Protein family information listed in the Pfam database[49]. <sup>3</sup>Promoter/sigma factor binding regions found on EcoCyc[39] based on computational predictions[50].

In general, key innovative mutations could be categorized as regulatory (R) or structural (S) (Table 1). Of the fifteen mutation events outlined in Table 1, eleven were categorized as regulatory (observed in all five successful substrate conditions) and four were categorized as structural (three of five successful substrate conditions). For D-lyxose, D-2-deoxyribose, and m-tartrate evolution experiments, mutations were observed within the coding regions of the predicted genes, namely yihS, rbsK, and dmlA (Table 1, Figs. S3-S5). Regulatory mutations, occurring in transcriptional regulators or within intergenic regions–likely affecting sigma factor binding and transcription of the predicted gene target–were observed for D-lyxose, D-2-deoxyribose, m-tartrate, and monomethyl succinate (Table 1). Observing more regulatory mutations is broadly consistent with previous reports[10, 24]. The regulatory mutations were believed to increase the expression of the target enzyme, thereby increasing the dose of the typically low-level side activity[18]. This observation is consistent with ‘gene sharing’ models of promiscuity and adaptation where diverging mutations that alter enzyme specificity are not necessary to acquire the growth innovation[18, 25]. Furthermore, although enzyme dosage could also be increased through duplication of genomic segments, this scenario was not commonly observed during the innovation phase of our experiments. Two large duplication events (containing 165 genes *(yqiG-yhcE*) and 262 genes *(yhiS-rbsK*), respectively) were observed only in the innovation phase adaption for growth on D-2-deoxyribose, and these regions did include the *rbsK* gene with the under-ground activity predicted to support growth (Table 1).

To identify the mutations that were causally involved in the nutrient utilization phenotypes, we re-introduced each key mutation (Table 1) into the ancestor wild-type strain using the genome engineering method (pORTMAGE)[26]. Genome editing was performed for screening mutation causality[27] on all novel substrate conditions, except for monomethyl succinate, which only contained a single mutation (Table 1). Furthermore, the large duplication in the D-2-deoxyribose strain could not be reconstructed using this method due to the limitations of the method. Individual mutants were isolated after pORTMAGE reconstruction, and their growth was monitored on the innovative growth medium over the course of one week. The growth test revealed that single mutations were sufficient for growth on D-lyxose, D-arabinose, and m-tartrate, but with varying lengths of time for growth to be detected depending on the mutation present (Table S3). For example, in the case of D-lyxose, growth was detected in YihW Δ2bp mutant cells in approximately 3-4 days, compared to 5-7 days for YihS single mutation cells. Interestingly, in the case of D-2-deoxyribose, an individual mutation (either the N20Y rbsK or the ΔrbsR mutation) was not sufficient for growth, thereby suggesting that the mechanism of adaptation to this substrate is more complex, requiring multiple mutation events (in this case, both regulatory and structural mutations). Overall, these causality assessments support the notion that underground activities open short adaptive paths towards novel phenotypes.

Were the mutations observed in our laboratory experiment relevant for environmental adaptations in the wild? The N20Y sole mutation observed in the RbsK enzyme during the evolution on D-2-deoxyribose served as a case study. Previous work has found that predominantly intestinal and extraintestinal strains of *E. coli*, as well as some *Salmonella* species, can use D-2-deoxyribose as a sole carbon source as they possess a pathogenicity island containing the deoxyribokinase deoK[28, 29, 30], which shares a 36% sequence identity with *rbsK* (BLASTp [31] expect (E) value 4e-29). Specifically, four such reported pathogenic strains in this set (three *E. coli* and one *Salmonella*) [28, 29, 30] were shown to grow on D-2-deoxyribose and possess a deoxyribokinase (DeoK) with a tyrosine residue at the equivalent N20Y position (Fig. S4). This information suggests that the N20Y mutation may have improved the ribokinase underground activity of RbsK in the mutant strain evolved here on D-2-deoxyribose and enabled growth in this environment similar to the capabilities of the strains that possess the DeoK enzyme (see Fig. S5 for a structural comparison). Therefore, this case highlighted that the genetic basis of adaptation observed in the laboratory is indeed relevant to evolution in the wild.

### 2.3. Contribution of enzyme side activities to the optimization phase of adaptation

Once the roles of mutations acquired during the innovation phase were established, adaptive mechanisms required for optimizing or fine-tuning growth on the novel carbon sources were explored. Specifically, of major interest for this study was the role of enzyme promiscuity during this second ‘optimization’[11] phase of the evolutions. Analysis of mutations in the optimization phase led to identification of additional promiscuous enzyme activities, above and beyond the innovative mechanisms, impacting the phenotypes of the evolved strains in four of the five nutrient conditions (Table 2). Discovery of these optimizing activities was driven by a systems-level analysis consisting of mutation, enzyme activity, and transcriptome analyses coupled with computational modeling of optimized growth states on the novel carbon sources.

**Table 2:**
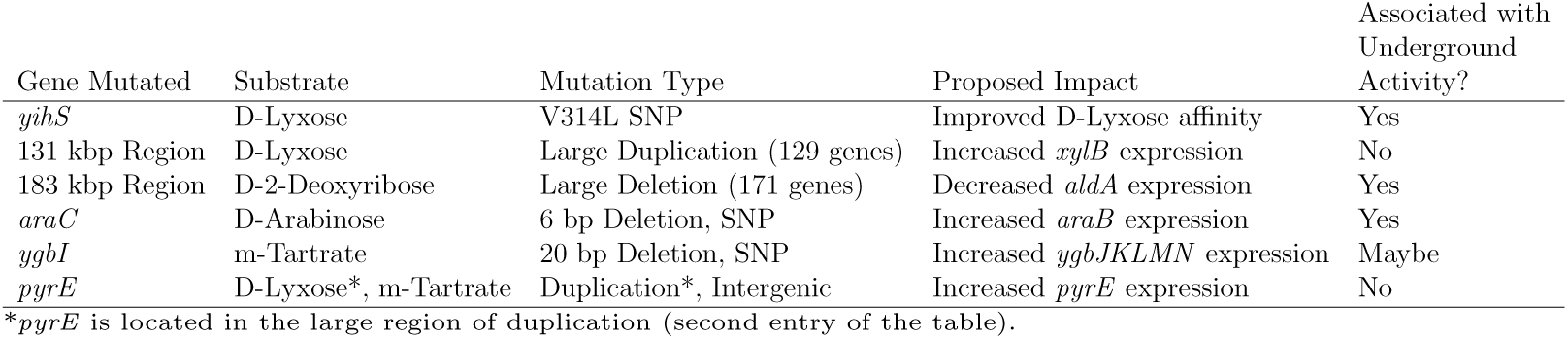
Optimizing Mutations.

The ‘optimization’ phase of the evolution experiments consisted of serially passing cultures in the early exponential phase of growth in order to select for cells with the highest growth rates (Fig. 1A). Marked and repeatable increases in growth rates on the non-native carbon sources was observed in as few as 180-420 generations (Table S1). Whole genome sequencing of clones was performed at each distinct growth-rate ‘jump’ or plateau during the optimization phase (see arrows in Fig. 1B, Fig. S1). Such plateaus represent regions where a causal mutation has fixed in a population[20]. Out of the total set of 41 mutations identified in the growth optimization regimes (Datasets S1, S2), a subset (Table 2) was explored. This subset consisted of genes which were repeatedly mutated in replicate experiments or across all endpoint sequencing data on a given carbon source. To unveil the potential mechanisms for improving growth on the non-native substrates, the transcriptome of initial and endpoint populations (right after the innovation phase and at the end of the optimization phase, respectively) was analyzed using RNAseq. Differentially expressed genes were compared to genes containing optimizing mutations (or their direct targets) and targeted gene deletion studies were performed. Additionally, for the D-lyxose experiments, enzyme activity was analyzed to determine the effect of a structural mutations acquired in a key enzyme during growth optimization.

Several mutations acquired during the optimization phase leading to large gains in fitness were directly linked to the influence of enzyme promiscuity. A clear example of optimizing mutations involved with optimization were those acquired during the D-lyxose experiments that were linked to enhancing the secondary activity of YihS, the enzyme also involved in the initial innovation. Protein structural mutations were observed beyond those observed during after the initial innovation. Structural mutations are believed to improve the enzyme side activity to achieve the optimization, and this effect was experimentally verified. The effects of structural mutations on enzyme activity were examined for the YihS isomerase enzyme that was mutated during the D-lyxose evolution (Fig. 1B, Table 1). The activities of the wild-type YihS and three mutant YihS enzymes (YihS R315S, YihS V314L + R315C, and YihS V314L + R315S) were tested in vitro. A cell-free in vitro transcription and translation system[32, 33] was used to express the enzymes and examine conversions of D-mannose to D-fructose (a primary activity[34]) and D-lyxose to D-xylulose (side activity) (Fig. 2A, Fig. S6). The ratios of the turnover rates of D-lyxose to the turnover rates of D-mannose were calculated and compared (Fig. 2B). The double mutant YihS enzymes showed approximately a ten-fold increase in turnover ratio of D-lyxose to D-mannose compared to wild type (P <0.0003, ANCOVA). These results suggest that the mutations indeed shifted the affinity towards the innovative substrate (enzyme side activity), while still retaining an overall preference for the primary substrate, D-mannose (ratio <1). This is in agreement with ‘weak trade-off’ theories of the evolvability of promiscuous functions[2] in that only a small number of mutations could result in significant improvements in the promiscuous activity of an enzyme without greatly affecting the primary activity.

**Figure 2:**
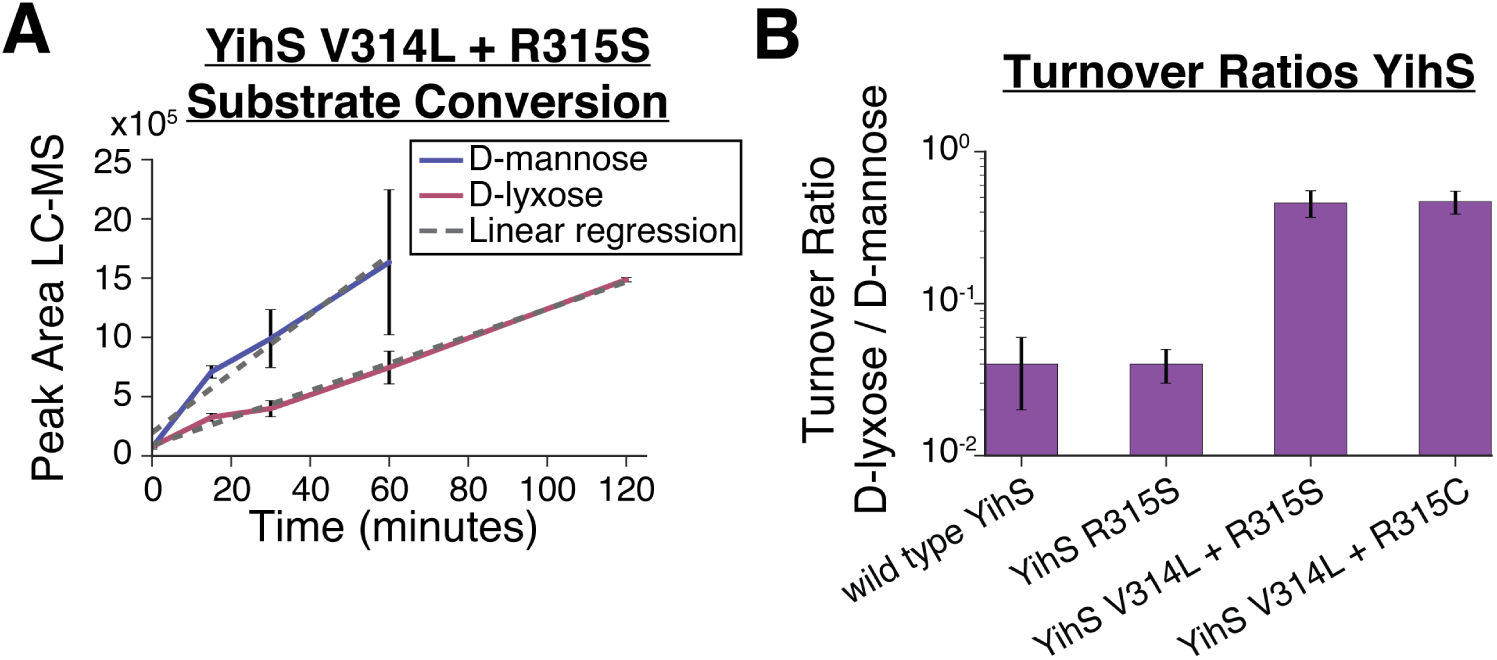
Evaluation of enzymatic activity for the wild-type and mutated promiscuous enzyme, YihS. A) YihS V314L + R315S mutant enzyme activity on D-mannose and D-lyxose. LC-MS was used to analyze YihS activity at saturating substrate concentrations to compare turnover rates on each substrate. Product formation was followed over time at a constant enzyme concentration. Turnover rates were calculated using linear regression. B) Turnover ratios of substrate conversion of D-lyxose / D-mannose are shown for the wild type YihS and mutant YihS enzymes. A ratio <1 indicates a higher turnover rate on D-mannose compared to D-lyxose. Error bars represent standard error calculated from the linear regression analysis.

Another clear example of an important optimizing mutation was found in the D-arabinose experiments occurring in the araC gene, a DNA-binding transcriptional regulator that regulates the araBAD operon involving genes associated with L-arabinose metabolism[35]. Based on structural analysis of AraC (Fig. 3A), the mutations observed in the two independent parallel experiments likely affect substrate binding regions given their proximity to a bound L-arabinose molecule (RCSB Protein Data Bank entry 2ARC)[36], possibly increasing its affinity for D-arabinose. Expression analysis revealed that the araBAD transcription unit associated with AraC regulation[37] was the most highly up-regulated set of genes (expression fold increase ranging from approximately 45-65X for Exp 1 and 140-200X for Exp 2, P <10^-4^) in both experiments (Fig. 3B). Further examination of these up-regulated genes revealed that the ribulokinase (AraB) has a similar kcat on four 2-ketopentoses (D/L-ribulose and D/L-xylulose)[38] despite the fact that araB is consistently annotated to only act on L-ribulose (EcoCyc)[39] or L-ribulose and D/L-xylulose (BiGG Models)[40]. It was thus reasoned that AraB was catalyzing the conversion of D-ribulose to D-ribulose 5-phosphate in an alternate pathway for metabolizing D-arabinose (Fig. 3C) and this was further explored.

**Figure 3:**
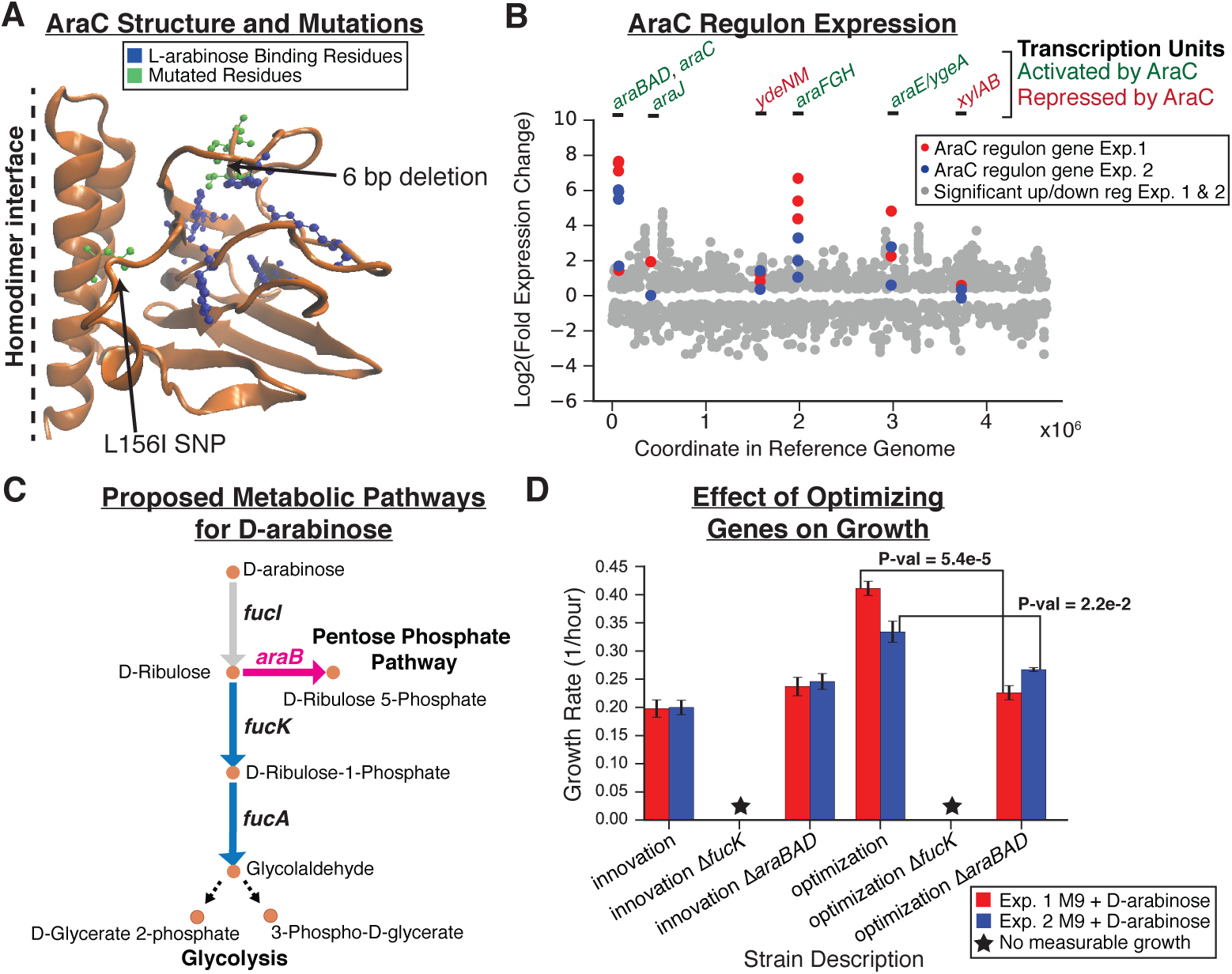
Optimization mutation analysis for D-arabinose evolution experiments. A) Structural mutations observed in sequencing data of Experiments (Exp.) 1 and 2 (green) as well as residues previously identified as important for binding L-arabinose (blue) are highlighted on one chain of the AraC homodimer protein structure. The six base pair deletion observed in Exp. 1 appears to be most clearly linked to affecting substrate binding. B) Expression data (RNAseq) for significantly differentially expressed genes (q-value <0.05). Scatter plot shows log2(fold change) of gene expression data comparing endpoint to initial populations for Exp. 1 and Exp. 2 (grey dots) with the location of the gene in the reference genome as the x-axis. Those genes that are associated with AraC transcription units are highlighted (red dots for Exp. 1 and blue dots for Exp. 2). Above the plot, the transcription units are labeled green if AraC activates expression (in the presence of arabinose) or red if AraC represses expression of those genes. C) The proposed two pathways for metabolizing D-arabinose. The pink pathway is enabled by the optimizing mutations observed in *araC*. D) Growth rate analysis of various innovation (starting point of optimization phase) and optimization (endpoint of optimization phase) strains with or without *fucK* or *araB* genes knocked out. Strains were grown on M9 minimal media with D-arabinose as the sole carbon source.

The role of the proposed second pathway in optimizing growth on D-arabinose was analyzed both computationally and experimentally. A flux balance analysis simulation of a model without the FucK associated ribulokinase reaction (the pathway of D-arabinose metabolism associated with innovative mutations), but with a non-zero flux through the AraB underground reaction, predicted in an approximately 10% higher simulated growth rate compared to when AraB is inactive (Fig. S7). This signaled the possibility of a growth advantage for using the *araB* enabled pathway and thus was explored experimentally. Experimental growth rate measurements of clones carrying either the *fucK* or *araBAD* genes knockouts showed that the FucK enzyme activity was essential for growth on D-arabinose for all strains analyzed (innovative and optimized) (Fig. 3D, Table S4). However, removal of *araB* from optimized endpoint strains reduced the growth rate of the strain to the approximate growth rate of the initial innovative strain (Fig. 3D), suggesting that the proposed *araB* encoded pathway (Fig. 3C) was responsible for enhancing the growth rate and therefore qualifies as fitness optimization. Putting this in the context of previous work, a similar pathway has been described in mutant *Klebsiella aerogens* W70 strains[41]. In the 1977 study, it was suggested that the D-ribulose-5-phosphate pathway (i.e., the *araB* pathway) is more efficient for metabolizing D-arabinose than the D-ribulose-1-phosphate pathway (i.e., the *fucK* pathway), possibly because the L-fucose enzymatic pathway requires that three enzymes recognize secondary substrates[41]. This conclusion supports the role of the optimization mutations observed here in *araC*. Overall, underground activities of both the *fuc* operon (innovative mutations) and *ara* operon (optimizing mutations) encoded enzymes were important for the adaptation to efficiently metabolize D-arabinose and the *ara* mutated operon did not solely support growth. Furthermore, computational analyses suggest that a similar mechanism of amplification of growth-enhancing promiscuous activities played a role in the m-tartrate optimization regime. Similarly, both independent evolutions on m-tartrate possessed a mutation in the predicted transcription factor, *ygbI*, with a resulting overexpression of a set of genes with likely promiscuous activity *(Supporting Text*, Fig. S8). Two additional proposed mechanisms for growth optimization on m-tartrate and D-lyxose were related to the primary activities of *pyrE* and *xylB* and are discussed in the *Supporting Text* (Fig. S8 and Fig. S10).

### 2.4. Loss of an enzyme side activity improves fitness

Analysis of the D-2-deoxyribose adaptation revealed a conceptually novel way by which alterations in promiscuous enzyme activities contribute to growth optimization. Several lines of observation suggested that suppression of a side reaction of aldehyde dehydrogenase A (AldA) enhanced growth on this novel carbon source. The optimizing mutation event in the D-2-deoxyribose evolution was a large deletion event spanning 171 genes (Fig. S9). Of these, the metabolic gene that was most significantly expressed in the ancestor (i.e., before the large deletion) was *aldA* (Fig. 4A). AldA has been described as a broad substrate specificity enzyme and has shown catalytic activity on acetaldehyde[42]. Turning to computational modeling to understand the impact of an active AldA, showed that forcing increased flux through acetaldehyde to acetate conversion decreased the overall growth rate (Fig. 4B, C; Dataset S3). Together, these findings indicate that the large deletion event observed in the D-2-deoxyribose endpoint selected against the AldA side activity, leading to improved growth. This scenario suggests that not only enhancement, but also suppression of side reactions can play a pivotal role in adaptation to novel environments.

**Figure 4:**
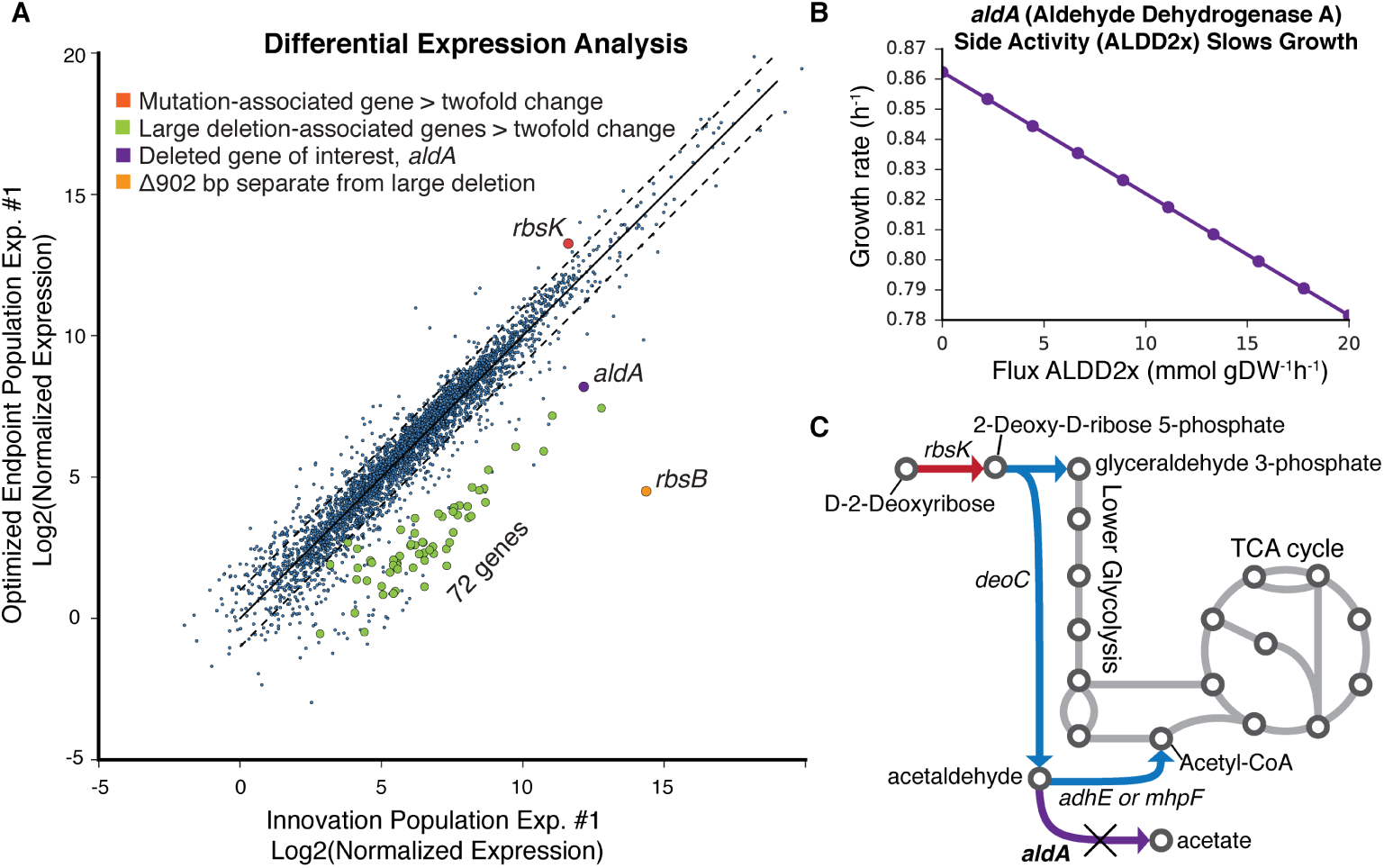
Optimization mutation analysis for the D-2-deoxyribose experiment. A) RNAseq expression data represented as log2(normalized expression) initial population samples compared to the endpoint population sample for experiment (Exp.) 1. Highlighted in red is *rbsK* associated with small mutation events and in green are genes associated with the large deletion. The *aldA* gene is highlighted in purple as a more highly expressed gene of interest that was within the large deletion region in the optimized endpoint population. The deleted genes have non-zero expression values in the optimized endpoint population, which can be explained by evidence that a fraction of the population does not contain the deletion (Fig. S9). B) A flux balance analysis plot showing the effect of flux through the reaction associated with *aldA* on growth rate. Flux through this reaction is predicted to have a negative effect on growth rate. C) A pathway map highlighting predicted pathways for metabolizing D-2-deoxyribose. Starting with D-2-deoxyribose in the upper left, the first reaction is catalyzed by the enzyme associated with the *rbsK* gene noted in red as it was a key gene mutated in the initial innovation population. The following reactions in blue are predicted to feed into lower glycolysis and the TCA cycle. The aldA-associated unfavorable underground reaction, converting acetaldehyde to acetate is highlighted in purple and marked with an X to note its deletion in an optimized endpoint population.

## 3. Conclusions

Taken together, the results of this combined computational analysis and laboratory evolution study show that enzyme promiscuity is prevalent in metabolism and plays a major role in both phenotypic innovation and optimization during adaptation. It was demonstrated that enzyme side activities can confer a fitness benefit in two distinct ways. First, side activities contributed to the establishment of novel metabolic routes that enabled or improved the utilization of a new nutrient source. Second, suppression of an undesirable underground activity that diverted flux from a newly established pathway conferred a fitness benefit.

The results of this study have direct relevance for understanding the role of promiscuous enzymatic activities in evolution and for utilizing computational models to predict the trajectory and outcome of molecular evolution[14, 43]. Here, it was demonstrated that computational metabolic network models which include the repertoire of enzyme side activities made it possible to predict the genetic basis of adaptation to novel carbon sources. As such, systems models and analyses are likely to contribute significantly towards representing the complex implications of promiscuity in theoretical models of molecular evolution[43]. Furthermore, the evolution of new gene functions from secondary promiscuous activities has been proposed by multiple models assuming functional gene divergence from a common ancestor following gene duplication events[44, 45, 46, 7, 47, 48] and the findings and strains from this study are relevant towards better understanding such models. Finally, the computational and subsequent approaches developed in this work can be leveraged to understand promiscuous activity in engineered strains for industrial biotechnology and in the adaptation of pathogenic microbes.

## 4. Materials and Methods

Flux balance analysis and sampling *in silico* methods utilized in this work are described in *SI Materials and Methods*. Detailed information regarding the laboratory evolution experiments, growth media composition, and whole genome sequencing and mutation analysis is provided in *SI Materials and Methods*. Furthermore, details regarding the pORTMAGE library construction and mutant isolation as well as the cell-free in vitro transcription, translation enzyme activity characterizations performed are also provided in *SI Materials and Methods*. Experimental methods utilized to analyze optimization regime mechanisms of adaptation including RNA sequencing, phage transduction mutagenesis, and individual mutant growth characterizations are included in *SI Materials and Methods*. The RNAseq data is available in the Gene Expression Omnibus (GEO) database under the accession number GSE114358. Data analysis, computation, and statistical analysis, unless otherwise specified in *SI Materials and Methods*, were conducted using the scientific computing Python library SciPy (http://www.scipy.org/) in a Jupyter Notebook (http://jupyter.org/).

## 5. Acknowledgments

We would like to thank Richard Szubin and Ying Hefner for assistance with strain resequencing and P1-phage transduction mutagenesis, as well as Elizabeth Brunk for helpful discussion. We also thank Suzanne Kosina for her assistance with LC-MS experiments. We also thank Samira Dahesh and the Nizet Lab for their assistance with Bioscreen experiments. This work was partially supported by The Novo Nordisk Foundation Grant Number NNF10CC1016517. TRN and MDR acknowledge funding from ENIGMA-Ecosystems and Networks Integrated with Genes and Molecular Assemblies (http://enigma.lbl.gov), a Scientific Focus Area Program at Lawrence Berkeley National Laboratory is based upon work supported by the U.S. Department of Energy, Office of Science, Office of Biological & Environmental Research under contract number DE-AC02-05CH11231. AN was supported by a PhD fellowship from the Boehringer Ingelheim Fonds. CP and BP acknowledge funding from the ‘Lendlet’ Programme of the Hungarian Academy of Sciences, The Wellcome Trust and GINOP-2.3.2-15-2016-00014 (EVOMER). CP was also supported by the European Research Council (H2020-ERC-2014-CoG).

## 6. Author Contributions

GIG, TES, RAL, TRN, RAN, CP, BP, BOP, and AMF designed the research; GIG, TES, RAL, ZAK, AN, HP, and MdR performed research; GIG, TES, ZAK, AN, MdR, BP, and AMF analyzed data; and GIG, BP, and AMF wrote the paper.

